# Microstimulation of primate neocortex targeting striosomes induces negative decision-making

**DOI:** 10.1101/668194

**Authors:** Satoko Amemori, Ken-ichi Amemori, Tomoko Yoshida, Georgios K. Papageorgiou, Rui Xu, Hideki Shimazu, Robert Desimone, Ann M. Graybiel

## Abstract

Affective judgment and decision-making are strongly modulated by the pregenual anterior cingulate (pACC) and caudal orbitofrontal (cOFC) cortical regions. By combining MRI-guided electrical microstimulation with viral tracing methods in non-human primates, we demonstrate that circumscribed pACC and cOFC microstimulation sites that induce negative decision-making preferentially project to striosomes in the anterior striatum. These results outline a behaviorally important circuit from pACC/cOFC to striosomes causally modulating decision-making under emotional conflict.

---

Microstimulation applied to the neocortex has been of great benefit in examining sensorimotor systems, but applying such methods to cortical regions related to the limbic system and emotion-related performance has been more difficult, both because the behaviors examined are complex and because these cortical regions have not been fully mapped. Few studies have combined microsimulation and behavioral analysis with anatomical output tracing of the stimulated regions. This triple analysis is crucial in order to delineate functional circuits related to affect. Here we developed a method for such triple circuit analysis. We chose to stimulate the pACC, as this region has been shown by microstimulation to affect motivationally challenging cost-benefit decision-making and to project to the striatum^1,2^. However, there is no evidence definitively identifying a corticostriatal circuit related to such challenging decision-making, known in humans to be deleteriously affected in neuropsychiatric disorders including anxiety and depression^3^. The presumed rodent homologue of the pACC has been implicated in circuits related to the striosome compartment of the striatum^4–6^, a highly distinct set of labyrinthine zones (‘striosomes’) distinguished from the surrounding matrix by their neurochemical composition^7,8^ and connections with the limbic system^9,10^. Potential homologies between rodent and primate, however, have been questioned, especially for medial prefrontal corticostriatal circuit^11^. Examining these cortico-striosomal circuits in non-human primates, therefore, has special value^12^, as their decision-related circuitry is considered to be more nearly homologous to cortical regions that in humans^13^ are implicated in neuropsychiatric disorders^3^.

We have here addressed this need in three ways. First, we applied cortical microstimulation to identify cortical regions that affected cost-benefit decisions in an approach-avoidance (Ap-Av) task used in humans to differentiate anxiety and depression^1^. We combined this with virus-based anatomical identification of the corticostriatal projections of the behaviorally defined cortical regions. In addition, we explored with this behaviorally identified microstimulation-targeting viral tracing to the cOFC, which is connected to the pACC and limbic regions and putatively with striosomes in anterior striatum^2,14^. Finally, we used fMRI to determine how such cortical microstimulation affected blood oxygen level dependent (BOLD) activity in the striatum to determine whether microstimulation of the behaviorally determined sites indeed activated the striatum. Throughout, we focused on the pACC, known to influence Ap-Av decision-making toward avoidance^1^, and the cOFC, not so far explored in this task-setting.

Three monkeys, S, Y, and P, learned to perform the Ap-Av decision-making task^1^ (Fig. 1a; Methods). After viewing combined stimuli indicating relative amounts of reward (juice) and punishment (airpuff to face), the monkeys indicated their choices by joystick movements to accept or reject the offer. Guided by MRI, we lowered electrodes to circumscribed locations in the pACC and cOFC to test whether microstimulation induced a significant change in the decision-making patterns (Supplementary Fig. 1a-c). We examined the effects of focal microstimulation (200-μs pulses delivered at 200 Hz, 100-200 μA) applied at 43 pACC and 35 cOFC sites during the first 1.0 s of the 1.5 s cue period. We compared the decision-making patterns between blocks of trials without stimulation (*Stim-off* block of 150-250 trials) and those with stimulation (*Stim-on* block of 150-250 trials) (Methods).

**Fig. 1.**
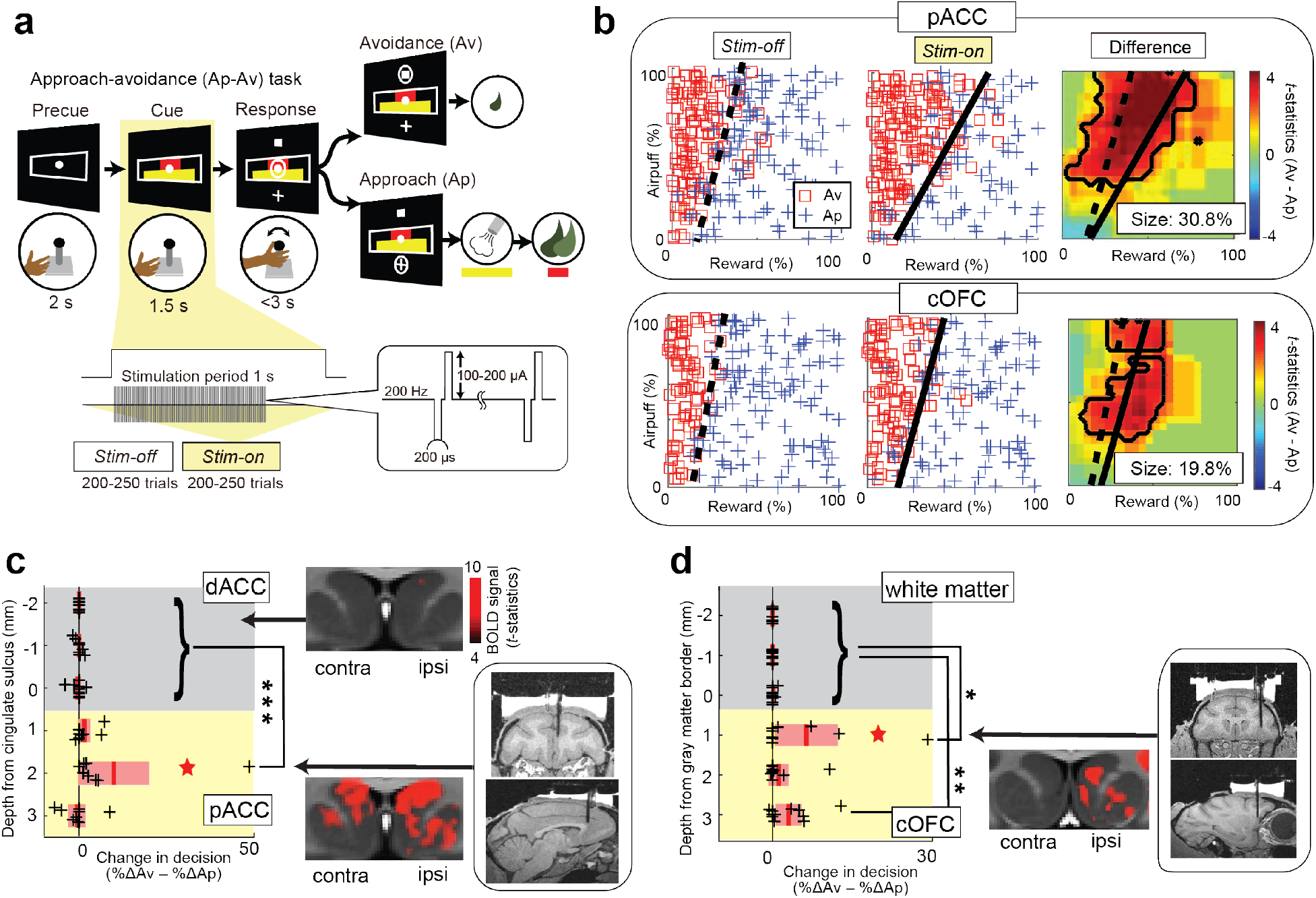
Task and Behavior. **a,** Approach-avoidance (Ap-Av) task (see Methods). **b,** Example of decision pattern affected by pACC (top) and cOFC (bottom) microstimulation, illustrated by choice: before (*Stim-off*, left) and during (*Stim-on*, middle) stimulation (*Stim-off*, left), and smoothed decision difference between the two blocks (right, t-statistics). Dashed and solid lines indicate decision boundary. **c,** pACC microstimulation increased Av choices compared to dACC stimulation (Wilcoxon signed rank test, \*\*\**P* < 0.001). Left: Size of increase in Av subtracted by that in Ap (black cross: %ΔAv – %ΔAp) plotted for the depth from cingulate sulcus. Gray, dACC. Yellow, pACC. Red star, experiment in **b**. Red lines, mean. Pink bars, standard errors of the means (SEM). Middle: Striatal fMRI signal induced by dACC (top) and pACC (bottom) stimulation. BOLD signal intensity over threshold (t-statistics = 4) shown by the right color bar. Right: Viral injection sites determined by the stimulation procedure. **d,** cOFC microstimulation increased Av choices; conventions as in **c**. The size of increase in Av induced by stimulation at two sites in cOFC gray matter was significantly larger than that in the white matter (\**P* < 0.05, \*\**P* < 0.01). Middle: Striatal fMRI signal induced by ipsilateral cOFC stimulation.

We compared the sizes of changes in decision induced by the microstimulation in the ventral bank of the anterior cingulate sulcus (pACC) to those in the dorsal sulcal bank (dACC) in monkeys S, Y and P. We binned those stimulation effects for every 1 mm from the cingulate sulcus and found that the size of increase in Av induced by the pACC stimulation 1 mm below the sulcus was significantly greater than that induced by the dACC stimulation (two-sided signed rank test, *P* < 0.001), confirming our previous evidence^1^ (Fig. 1b,c and Supplementary Fig. 2). In the cOFC of monkeys Y and P, we compared microstimulation effects between white and gray matter by binning the effects for every 1 mm from the gray matter border, and found two cOFC sites at which the microstimulation induced significant increase in Av choices compared to those in the dorsally adjoining white matter (signed rank test, *P* < 0.05) (Fig. 1b,d and Supplementary Fig. 3). These effects in the cOFC have not been reported before, and suggest a second cortical source of negative bias in decision-making.

To examine whether stimulation of the pACC and cOFC actually induced striatal activation, we delivered microstimulation in monkey N (anesthetized) during periods in which we performed fMRI imaging^15,16^ (Methods), targeting the microstimulation at the cOFC region corresponding to the locations at which stimulation increased avoidance in monkeys S, Y and P. Control dACC microstimulation did not induce significant striatal BOLD activation, whereas the targeted pACC stimulation activated the striatum bilaterally. In the cOFC, microstimulation primarily activated striatal regions ipsilateral to the stimulation sites (Fig. 1c,d and Supplementary Fig. 4). Thus stimulation of the behaviorally effective sites, but not nearby sites in the pACC and cOFC, indeed activated the striatum.

We next injected viral tracers (AAVDJ-CMV-hrGFP or AAVDJ-CMV-mCherry) at these behaviorally identified pACC and cOFC sites (Supplementary Table; Figs. 2c, 3c and Supplementary Fig. 1d-f; Methods). Seven weeks later, we performed dual immunostaining of corticostriatal afferents for hrGFP or mCherry antibody (virus tracer) and KChIP1 (striosome marker) on the same striatal sections. For the pACC, the injection sites in monkeys S and Y were pre-identified by the stimulation results of the monkeys, and the virus-infected regions around the injection sites of monkeys A and N were confirmed to include the locations of stimulation-effective sites in the post-mortem examination (Fig. 2c). Behaviorally effective microstimulation sites in the pACC exhibited preferential targeting of subsets of striosomes, mainly in the anterior striatum (Fig. 2a and Supplementary Fig. 5). For the cOFC, we predetermined the injection sites of monkeys A and N by the effective sites of the microstimulation experiments performed in monkeys Y and P, and we confirmed histologically that the infected regions around the injection sites included the locations of the effective sites (Fig. 3c). The effective site projections to the striatum preferentially labeled subsets of striosomes in more ventral regions of the striatum (Fig. 3a and Supplementary Fig. 6). Thus there was a significant striosomal bias in corticostriatal projections from the pACC and cOFC sites inducing increased avoidant behavior.

**Fig. 2.**
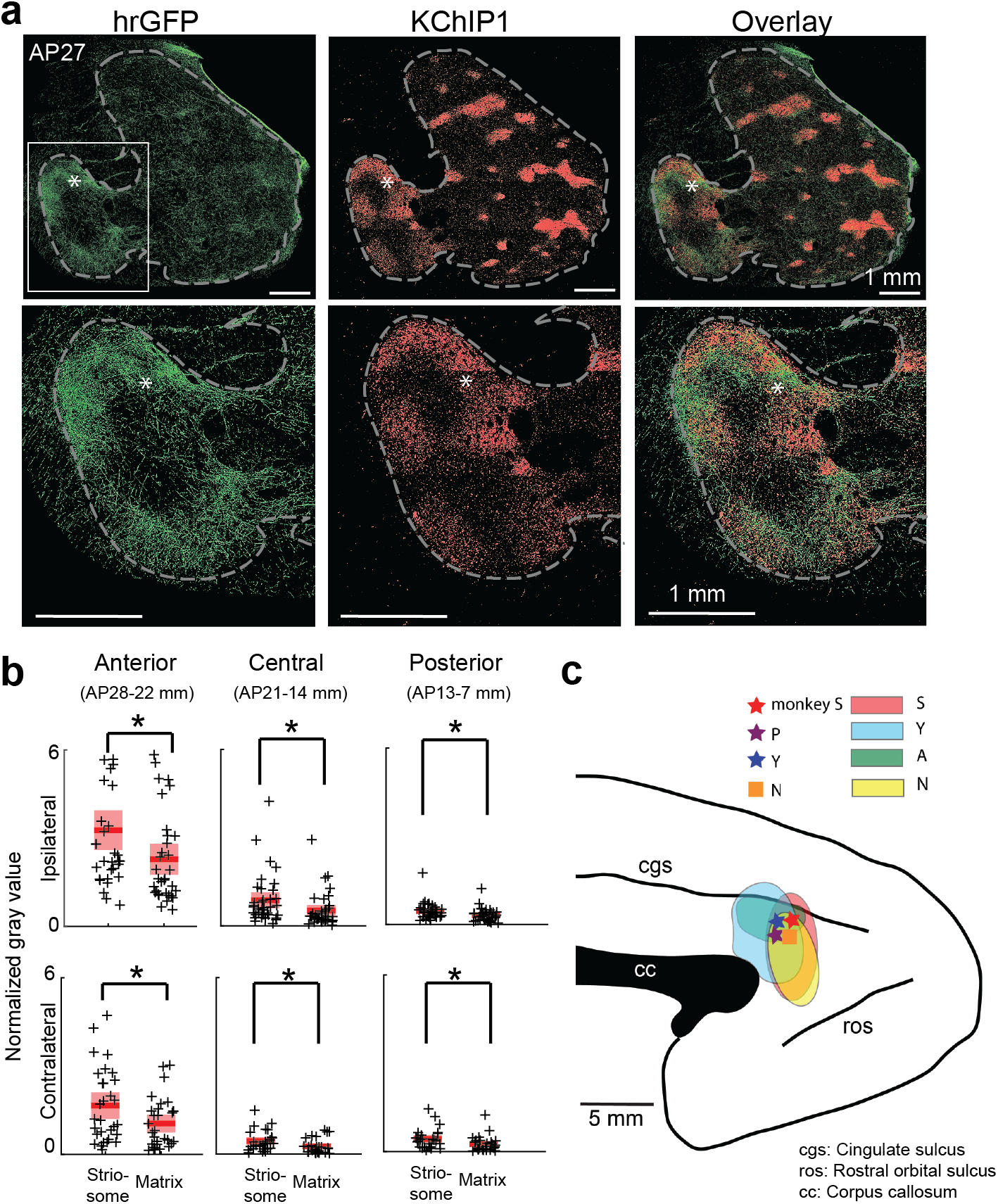
pACC projections to striosomes. **a,** pACC projection (hrGFP, green, left), KChIP1-positive striosomes (red, middle) and merged images (right), with enlarged views of the anterior striatum (framed by dashed line) at bottom. Asterisks indicate the same positions. **b,** Quantification of pACC projections to striatum. Tracer density (cross) was represented by normalized gray value of each striatal region and was calculated separately for striosomes and matrix (Methods), and for anterior, central and posterior regions. Red bars, means. Pink bars, SEM. Tracer densities were significantly greater in striosomes than in matrix in both hemispheres (paired *t*-test, \**P* < 0.05). **c,** Effective microstimulation and viral injection sites (stars) in monkeys S (red), P (purple) and Y (blue), and stimulation site in monkey N during fMRI (square). Colored regions, extents of viral injection sites (S, red; Y, blue; A, green; N, yellow).

**Fig. 3.**
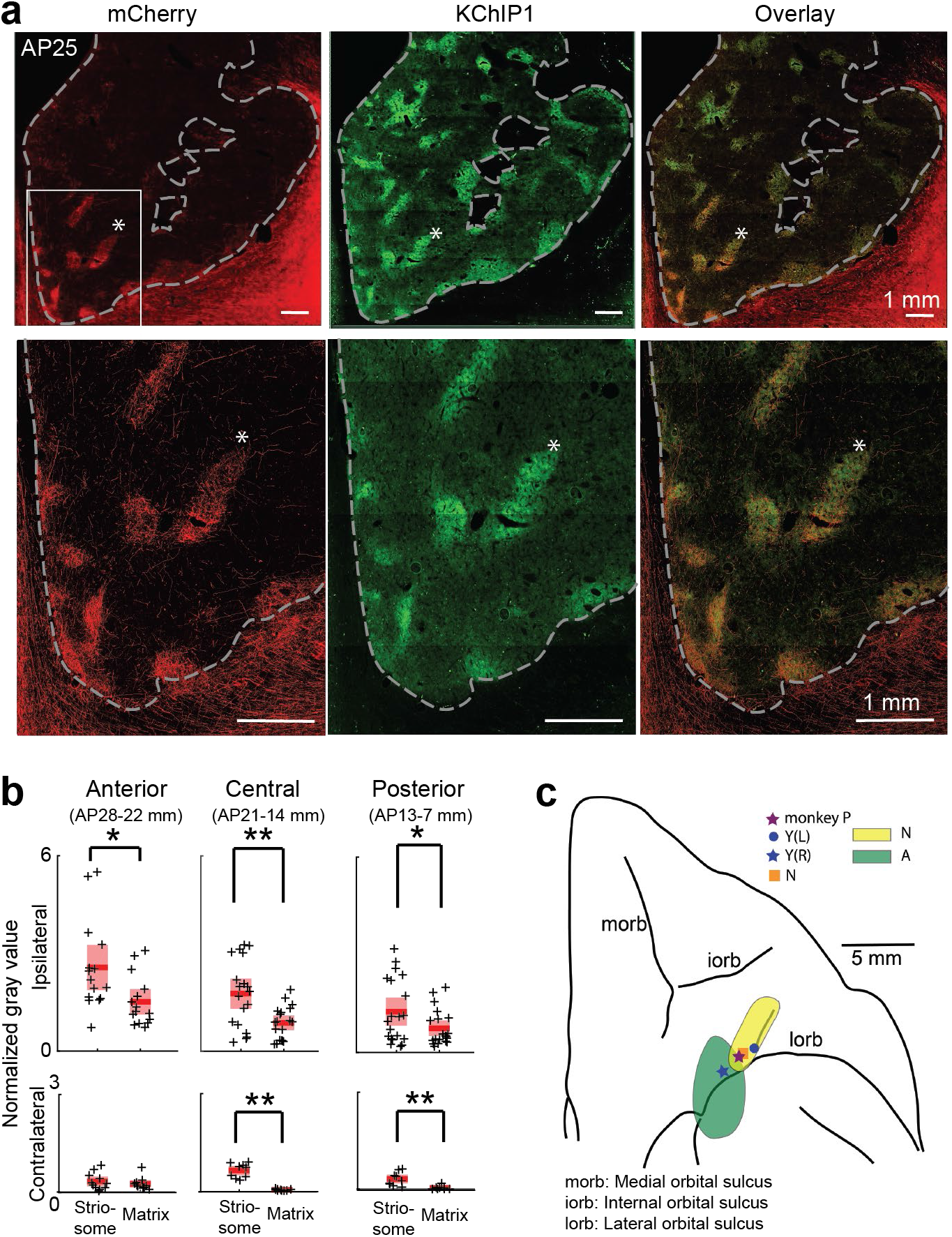
cOFC projections to striosomes. Conventions as in Fig. 2. **a,** cOFC projection (mCherry, red, left), KChIP1-positive striosomes (green, middle) and merged images (right). **b,** cOFC projections to striosomes were significantly different from those to matrix (\**P* < 0.05, \*\**P* < 0.01). **c,** Effective microstimulation sites and extents of viral injection sites.

We quantitatively estimated for the sites producing increased avoidance the virally labelled zones in the striosomes and matrix in extensive anterior, central, and posterior striatal regions (Figs. 2b, 3b and Supplementary Figs. 7-8; Methods). The densities were significantly higher in striosomes than in matrix (paired t-test, *P* < 0.05 for ipsilateral striatum), suggesting that pACC and cOFC sites at which microstimulation could alter conflict decision-making both provided relatively concentrated striatal input to striosomes. As is commonly found, even concentrated projection zones are accompanied by weaker staining beyond their borders^14^. That was the case here. As a further control, we therefore injected AAVDJ-CMV-mCherry (Supplementary Table) into the caudal ventral cingulate motor area in monkey I, posterior to the effective pACC zone. The matrix compartment, but not striosomes, was mainly labelled (Supplementary Fig. 9). These results indicated a decline of the striosome-projecting cingulate cortex posterior to the pACC, and suggest that only a limited region of the cingulate cortex sends signals to the anterior striosomal system.

These findings are the first to demonstrate that cortical sites in non-human primates behaviorally identified as modulating conflict decision-making send corticostriatal innervations favoring the striosome compartment over the surrounding matrix. The sites at which microstimulation increased avoidance preferentially targeted striosomes in each monkey, in at least some topographically distinct innervation zone. The pACC and cOFC are interconnected and related to limbic regions^14^. Our results in non-human primates provide causal evidence that circumscribed, behaviorally identified zones in the pACC and cOFC participate in corticostriatal circuits that contribute critically to negative decision-making under challenging cost-benefit decision-making. Identifying regions with corticostriatal input differentially recruited in approach choices, or both approach and avoidance, is an important future goal^17^. Differential effects of neurodegenerative disorders on striosomes and matrix have been indicated by post-mortem studies in humans^18–20^. The identification here of behaviorally identified cortical sites with differential striosome-matrix connectivity could help to unravel the circuit-level basis affected in such clinical disorders.

## Acknowledgements

We thank Drs. Helen Schwerdt, Simon Hong, Leif Gibb and Patrick Tierney for help with experiments; Margo Cantor, Jonathan Gill, Caitlin Erickson, Lauren Stanwicks, Alexandra Burgess and Polly Weigand for help with monkey care and training; Jannifer Lee, Michael Riad, Drs. Christian Wuethrich and Thomas Diefenbach for help with histology and imaging; Hanna May, Zahra Malik and Emily Chung for help with histology and image analysis; Henry Hall and Dr. Dan Hu for help with many aspects of this work; and Dr. Yasuo Kubota and Sarah Ryan for help with manuscript preparation. We thank Athinoula A. Martinos Imaging Center at the McGovern Institute for Brain Research, MIT for use of MRI; and Imaging Core at the Ragon Institute of MGH, MIT, and Harvard for use of TissueFAXS scanner. This research was supported by the National Institutes of Health (R01 NS025529), the CHDI Foundation (A-5552), the Office of Naval Research (N00014-07-1-0903), the Army Research Office (W911NF-16-1-0474), MEXT KAKENHI (18H04943 and 18H05131), the Simons Center for the Social Brain, the Saks Kavanaugh Foundation, the Naito Foundation, the Uehara Foundation, and Amy Sommer and Judy Goldberg.

## Author contributions

S.A., K.A., and A.M.G. designed the experiments. S.A., K.A., H.S. and A.M.G. performed the surgeries and injection experiments. S.A. and K.A. collected the recording and stimulation data and analyzed the electrophysiological data with critical feedbacks from A.M.G. S.A., T.Y. and A.M.G. performed histology and analyzed the anatomical data. S.A., G.P., R.X. and R.D. performed fMRI experiment, and G.P. analyzed the fMRI data. S.A., K.A. and A.M.G. wrote the manuscript.

## Competing interests

The authors declare no competing interests.

## Data availability

The data generated and analyzed and codes used in this study are available from the corresponding author on reasonable request.References

## Methods

### Subjects and experimental conditions

Four female (S: 7.5 kg, P: 6.3 kg, Y: 5.8 kg, A: 6.5 kg) and two male (N: 13.5 kg, I: 13.0 kg) *Macaca mulatta* monkeys were used in microstimulation and/or viral injection experiments. All experimental procedures were approved by the Committee on Animal Care at the Massachusetts Institute of Technology and were in accordance with the Guide for Care and Use of Laboratory Animals of the United States National Research Council. Before training on behavioral tasks, monkeys S, P and Y were habituated to sitting in a monkey chair and to wearing a head-fixation device^21^. Monkey S was used in neuronal recording, microstimulation and virus injection experiments. Monkey P was used in neuronal recording and microstimulation experiments. Monkey Y was used in microstimulation and virus injection experiments. Monkeys A, N and I were used for anatomy experiments. Monkey N was also used for fMRI imaging with microstimulation. We injected virus tracers in the pACC and/or cOFC of four monkeys (S, Y, A and N). We injected virus tracers in the ventral part of the cingulate motor area of monkey I as control for pACC injections. The details of the procedures that have been made in each monkey were summarized in Supplementary Table 1.

### Task procedures and microstimulation

Three monkeys (S, P and Y) were trained to perform the approach-avoidance (Ap-Av) task shown in Fig. 1a. The task started when the monkey put its hand on the designated position in front of the joystick. After a 1.5-s fixation (precue) period, two bars appeared on the screen as a compound visual cue. The lengths of the red and yellow bars indicated the offered amount of food and airpuff delivered after approach choice, respectively. After the 1.5-s cue period, two targets (cross and square) and an open circle appeared on the screen, and then the monkey could move the circle to choose one of the two targets by the joystick. Cross target was associated with approach (Ap) choice, and the square target was associated with avoidance (Av) choice. The locations of the two targets were randomized across trials. After an Ap decision, both airpuff and reward were delivered as offered. After an Av decision, the monkey did not receive the offered airpuff or food but had a small amount of food to maintain the motivation. When the monkey made commission or omission errors, the airpuff was delivered at the strength associated with the length of the yellow bar. After the behavioral training, a plastic recording chamber was implanted on the skull. We used MRI to identify the target region for implanting platinum-iridium electrodes (impedance 1-2 MΩ; FHC, ME) (Supplementary Fig. 1a,b). Single unit activity was also recorded from the cOFC while the monkeys performed the Ap-Av task (Supplementary Fig. 3c). The neuronal recording results for the ACC were published in our previous paper^1,22^.

During the microstimulation experiment, stimulation-off (*Stim-off*) and stimulation-on (*Stim-on*) blocks were alternated every 200-250 trials. At each trial in the *Stim-on* block, single monopolar stimulation was applied for 1.0 s starting at the onset of cue presentation. The stimulation train consisted of 200-μs pulses delivered at 200 Hz. Each pulse was biphasic and was balanced with the cathodal pulse leading the anodal pulse. The current magnitude was 100–200 μA (Fig. 1a). In each microstimulation experiment, we compared the choice pattern in the *Stim-off* and *Stim-on* blocks. For each block, we represented the size of the changes in Ap-Av decision by t-statistics that were calculated from a decision matrix. To make the decision matrix, we first convolved the choice in each trial by 30-by-30 point square-smoothing windows. After the spatial smoothing, each choice datum was stacked at each point in the 100-by-100 decision matrix. We then used two-sided Fisher’s exact test to detect statistical differences between the *Stim-off* and *Stim-on* blocks (*P* < 0.05). To calculate the significance level of the difference in decision matrices between the *Stim-off* and *Stim-on* blocks, we measured the total change in decision frequencies as the sum of the increase in Ap and that in Av choices (i.e., %ΔAp + %ΔAv). We set a change of 5% as the threshold to discriminate effective and non-effective sessions because in our previous study the false-positive rate to misclassify non-effective as effective was less than 5% with the threshold^23^ (Supplementary Figs. 2, 3). After we defined its significance level, we used the difference in positive and negative effects (i.e., %ΔAv – %ΔAp) to clarify the size and direction of each effect. We note that the sum of increases (%ΔAp + %ΔAv) was the same as and the difference (%ΔAv – %ΔAp) of the increases in all effective sessions as they exhibited only an increase in Av or Ap for each session.

### Injection of virus tracers into stimulation-effective sites

In order to inject virus tracers into the behaviorally effective site that was identified by microstimulation, we implanted a guide tube that was used in both microstimulation and injection experiments (Supplementary Fig. 1). The guide tube was made with fused silica tubing (TSP250350; OD: 350 μm, ID: 250 μm) and implanted in the pACC of the right hemisphere and the OFC of both hemispheres in monkey Y. Then we inserted a platinum-iridium electrode (impedance: 3 kΩ; FHC, ME) into the guide tube. The guide tube was held by a small custom-made plastic manipulator for MRI (N0-035-001, Narishige, Japan). The manipulator was attached on top of a custom-made plastic grid that had a hole (outer diameter: 1 mm) at every 1 mm. A guide tube was inserted through each hole. A stopper made from epoxy was attached on top of the electrode to make the tip of the electrode always extend 1-2 mm from the bottom of the guide tube (Supplementary Fig. 1 a,b). For the pACC implant, we advanced the guide tube with the electrode by the micromanipulator, targeting the cingulate sulcus. For the cOFC, we implanted the guide tube with the electrode, targeting the white matter above the cOFC. We performed this procedure while taking MRI images of the brain. Thus we could visually confirm the location of the tip of both pACC and cOFC electrodes before starting the first microstimulation experiment. Prior to each microstimulation experiment, we advanced the guide tube holding the electrode ~500 μm, and then the monkey started to perform the task. If we found a stimulation-effective site, we stopped the guide tube at that site. The electrode tip was again visualized by MRI. Thus we could confirm that the tip of the electrode was below the cingulate sulcus (Fig. 1c) and was in the cOFC (Fig. 1d) at the effective sites. In monkey S and Y, after finishing all microstimulation experiments, we removed the electrode but left the guide tube at the effective sites. We measured the length of the electrode and made 32-gauge injection needle that had a stopper exactly at the same position as the electrode in order to inject the virus tracer to the effective sites (Supplementary Fig. 1 c,d). The needle was attached to 10 Hamilton syringe (#80008; Hamilton, NV).

### Injection of virus tracers

Monkeys A and I had a craniotomy over the frontal cortex under sevoflurane anesthesia, and a slit was made on the dura. We estimated the target positions for the injection by MRI images as well as the relative position from the midline (for the pACC and the CMA) and the arcuate sulcus (for the cOFC). After the injection, the slit on the dura was sutured, an artificial dura (DuraGen; IntegraLifeSciences, NJ) was placed to cover the suture, and then the skin was sutured. Monkey N had a craniotomy, and a custom-made chamber was attached over the pACC and cOFC of the right hemisphere^24^. We estimated the injection position by MRI images and the grid coordination. We used a fused silica guide tube for fMRI in monkey N, and the injection was followed by fMRI.

### Injection of virus tracers

We used two different virus constructs as neural tracers: AAV-DJ-CMV-hrGFP (genomic titer: 1.10E+14 vg ml^-1^; Infectious titer: 2.50E+09 IU ml^-1^) and AAV-DJ-CMV-mCherry (genomic titer: 2.10E+14 vg ml^-1^; Infectious titer: 3.30E+10 IU ml^-1^; Stanford Vector Core, CA). The virus tracers and their amount injected are summarized in Supplementary Table 1. The virus tracer was pressure-injected by a 10-μl Hamilton syringe with a 32-gauge injection needle (#80008, Hamilton, NV). For monkeys A, N and I, after we punched the dura matter by a metal guide tube (27-gauge thin wall stainless steel needle), the needle was advanced to the target position with the control of a stereotactic arm. For monkeys S and Y, we used pre-implanted guide tubes. The injection needle was lowered through the guide tube and stopped at the targeting depth. We injected virus tracers at three different depths along the injection track. Each position was 0.5 mm apart for the pACC and 1 mm apart for the cOFC. The top position was the effective stimulation site in monkeys S and Y, but was estimated in monkeys A, N and I. The injection speed was 0.05 μl min^-1^ and was controlled by an injection pump (QSI No. 53311, Stoelting Co., IL). The pump and the syringe were held by a stereotactic arm that was attached on a stereotactic frame. After the injection, the guide tubes were removed immediately except for monkey Y, in which the guide tube was kept in place until perfusion and the virus tracer leaked along the guide tube track. Thus, in monkey Y, the virus was overexpressed above the stimulation effective site in the cOFC, which prevented data analysis. Anti-biotics were given for ten days after the injection for all monkeys. We waited at least seven weeks after the injection to examine the virus expressions in the striatum.

### Histology

We deeply anesthetized the monkeys with an overdose of sodium pentobarbital, and they were perfused with 0.9% saline followed by 4% paraformaldehyde in 0.1 M phosphate buffered saline (PBS). Brains were kept in 4% paraformaldehyde for 3 days, and then electrodes were withdrawn in monkeys S, P and Y. Brains were kept in 4% paraformaldehyde for only 1 day in monkeys A, N and I. The brains were separated into the left and right hemispheres, and striatal blocks were made. The blocks were stored in 25% glycerol in 0.1% sodium azide (Sigma, 438456) in 0.1 M phosphate buffer (PB) at 4°C until being frozen in dry ice on a sliding microtome and cut into 40-μm coronal sections. Sections were stored in 0.1% sodium azide in 0.1 M PB.

For immunofluorescent staining, sections were rinsed 3 times for 2 min in 0.01 M PBS containing 0.2% Triton X-100 (Tx) (PBS-Tx; Sigma-Aldrich, T8787) and then were pre-treated with 3% H_2_O_2_ in PBS-Tx for 10 min. Sections were rinsed 3 times for 2 min in PBS-Tx and incubated in tyramide signal amplification (TSA) blocking reagent (PerkinElmer, FP1012) in PBS-Tx (TSA-block) for 60 min. The striatum sections were incubated with primary antibody solutions containing rabbit anti-hrGFP (for hrGFP; 240141, Agilent Technologies) or rabbit anti-RFP (for mCherry; Rockland, 600-401-379) and mouse anti-KChIP1 (UC Davis/NIH Neuro Mab Facility, #75-003) in TSA-block for 24 hrs at 4°C. After primary incubation, the sections were rinsed 3 times for 2 min in PBS-Tx and then incubated for 1 hr in the secondary antibody solution containing goat anti-mouse Alexa Fluor 647 [1:400] (A21236, Invitrogen). After the sections were rinsed 3 times for 2 min in PBS-TX, the sections were incubated in anti-rabbit polymer HRP (GTX83399, GeneTex) solution and then rinsed 3 times for 2 min. The sections were incubated in TSA-block containing Streptavidin 488 [1:2000] (for hrGFP; Jackson Immuno Research, 016-540-084) or Streptavidin 546 [1:2000] (for mCherry; Life Technologies, S11225). We rinsed the sections 3 times for 2 min in 0.1M PB, mounted onto glass slides and then coverslipped with ProLong Antifade Reagent (Life Technologies, P36930). Sections were examined microscopically, and the striatal regions were imaged with an automatized slide scanner (TissueFAXS Whole Slide Scanner; TissueGnostics, Vienna, Austria) fitted with 10X objectives. Three fluorescence filter cubes (Alexa 488 for hrGFP, Texas Red for mCherry and Cy5 for KChIP1) were used to image the sections. The images were viewed by TissueFAXS viewer software (TissueGnostics, Vienna, Austria) and exported to tiff images.

### Image analysis

We picked hrGFP/mCherry and KChIP1 images of the same section every ~18 sections (every ~720 μm) from the start to the end of the striatum. Striosome and matrix masks were generated from the KChIP1 image by k-mean clustering function of Matlab (Mathworks, MA). The hrGFP/mCherry image was transformed into grayscale. In each image, the density of gray value (0-255) per pixel in striosomes and matrix was calculated using the masks. The density was statistically compared between striosomes and matrix by two-sided paired sample *t*-test.

### Visualizing corticostriatal pathways in fMRI

Monkey N underwent electrical microstimulation (EM) and fMRI under anesthesia inside a 3T MRI scanner (Siemens, Erlangen, Germany) using a saddle-shaped single-loop 12.7-cm receive coil. During the initial anesthesia by ketamine (1 mg kg^-1^), we implanted a fused silica guide tube and inserted a tungsten electrode (impedance: 1-3 kΩ; FHC, ME) through the guide tube to the pACC and cOFC. The grid coordination and target sites for the EM-fMRI were the same as those for the injections and confirmed by MRI before starting the EM-fMRI sessions. We maintained anesthesia by continuous intravenous administration of propofol (3-4 mg kg^-1^), at an infusion rate of 0.1-0.6 mg kg^-1^ min^-1^. Respiratory rate was assessed by continuous capnographic evaluation, and the body temperature of the animal was kept constant throughout the procedure by heated water blankets and was monitored at the intervals between scans. Probes for measuring blood oxygenation (SPO_2_) and heart rate were attached, and these measures were recorded continuously. Before starting functional scans, Ferumoxytol (Feraheme), an iron oxide-based contrast agent, was administered intravenously (8 mg kg^-1^), which substantially increases the signal-to-noise ratio of fMRI^25^. The scanning session lasted ~4 hr in total.

During the session, four functional runs of EM-fMRI were performed for each region of interest (ROI), using a standard block design and parameters of microstimulation similar to previous studies in this field^15,26,27^. Specifically, each functional run included eight 30-s EM blocks interleaved with nine 30-s rest blocks in which no stimulation was delivered. During EM blocks, 210-ms pulse trains were delivered at a rate of 1 Hz, which consisted of 70 biphasic current pulses. A single pulse was composed of a 200-μs negative phase, a 100-μs interval, and a 200-μs positive phase. Inter-pulse interval was set at 2.5 ms so that the frequency of microstimulation was 333 Hz. The amplitude of both negative and positive phases was set at 500 μA^15^, because we found (data not shown) that this parameter provides a superior contrast-to-noise ratio of EM-fMRI compared to lower current levels (e.g., 300 μA or lower), which given our scanning parameters might render some EM-fMRI activations difficult to detect. Stimulation pulses were delivered in a monopolar configuration with a computer-triggered pulse generator (DS8000; World Precision Instruments) connected to an isolator (A365; World Precision Instruments). The triggering of pulses was controlled and synchronized with MRI data collection using a desktop computer and custom Matlab code based on Psychtoolbox. Both stimulation and reference electrodes were connected to the isolator through custom-made inductor-based low-pass filters both on the patch-panel and inside the bore of the magnet of the scanner room to avoid contamination of the MR images as well as excessive heat from radio frequency noise.

MRI data were analyzed with Freesurfer/FsFast. A high-quality T1-weighted image collected before the EM-fMRI session was used as a template, and the functional data were co-registered to this template in a two-step fashion, using the T1-weighted image collected during the EM-fMRI session as an intermediate template (same session). Besides common processing steps including slice-timing correction and motion correction, field maps collected close to the functional images (between which there was little movement of the animal) were used to correct geometric distortion of functional data and thus improve its alignment with same-session T1 template. EM-fMRI activation maps (t-score) were computed in the native functional voxel space using FsFast’s GLM implementation for each ROI separately and were interpolated to the high-quality T1 template for visualization.

