## Supplementary Figures 1-9 for "Microstimulation of primate neocortex targeting striosomes induces negative decision-making"

### Supplementary Information

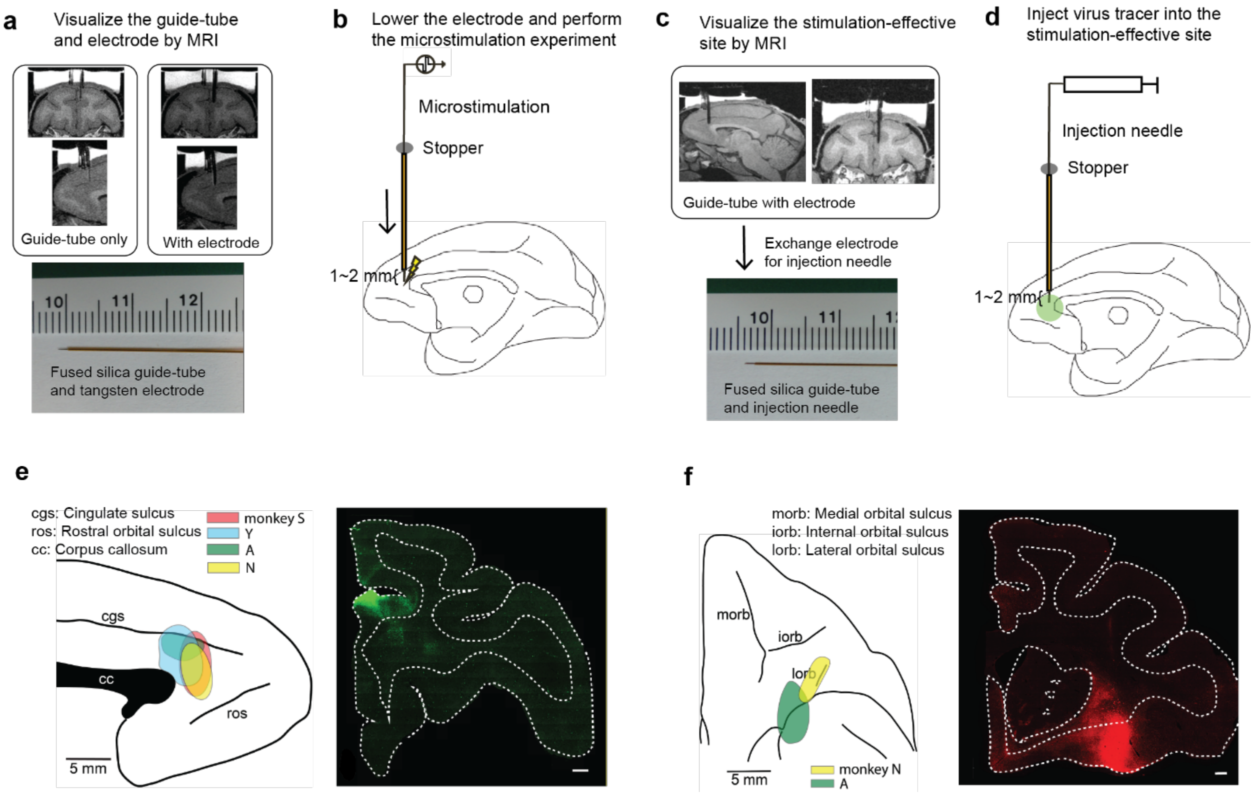


### Supplementary Figure 1

**Experimental design to deliver microstimulation and infuse virus tracers at the same target cortical site.**

**a)** We visualized the fused-silica guide tube lowered to the dACC region by MRI (top left panel). Then, we inserted a stimulation electrode through the guide tube until the stopper attached to the electrode prevented further lowering (top right panel). As we had already measured the distance between the tip of the electrode and that of the guide tube (bottom panel, 1-2 mm), we could estimate the position of the electrode tip and also estimate the artifact around the tip to be minimal (~1 mm). **b)** We lowered the guide tube and electrode, and performed the microstimulation experiment to map out the function of each zone in the dACC and pACC (Fig. 1b). We moved the guide tube and electrode every session toward the ventral part. **c)** After functional mapping, we moved the electrode tip to the effective site and again visually identified the site by the MRI with the electrode (top panel and Fig. 1c). To infuse the tracer virus, we left the guide tube and replaced the stimulation electrode with an injection needle. We had already put the stopper on the needle so that the needle tip would end up at the exact location where the electrode tip was (bottom panel). **d)** We infused the tracer virus to the stimulation-effective site via the needle inserted into the guide tube. For the cOFC, we predetermined the infusion site in two monkeys (A and N) based on the results of other monkeys (P and Y), but we later confirmed that they were at the corresponding sites by performing histology (Fig. 3c). **e-f)** Schematic diagram (left) and representative images (right) of tracer injection sites in pACC (e) and cOFC (f). Each color represents each monkey (red: S, blue: Y, green: A, yellow: N). Scale bars in images represent 1 mm.

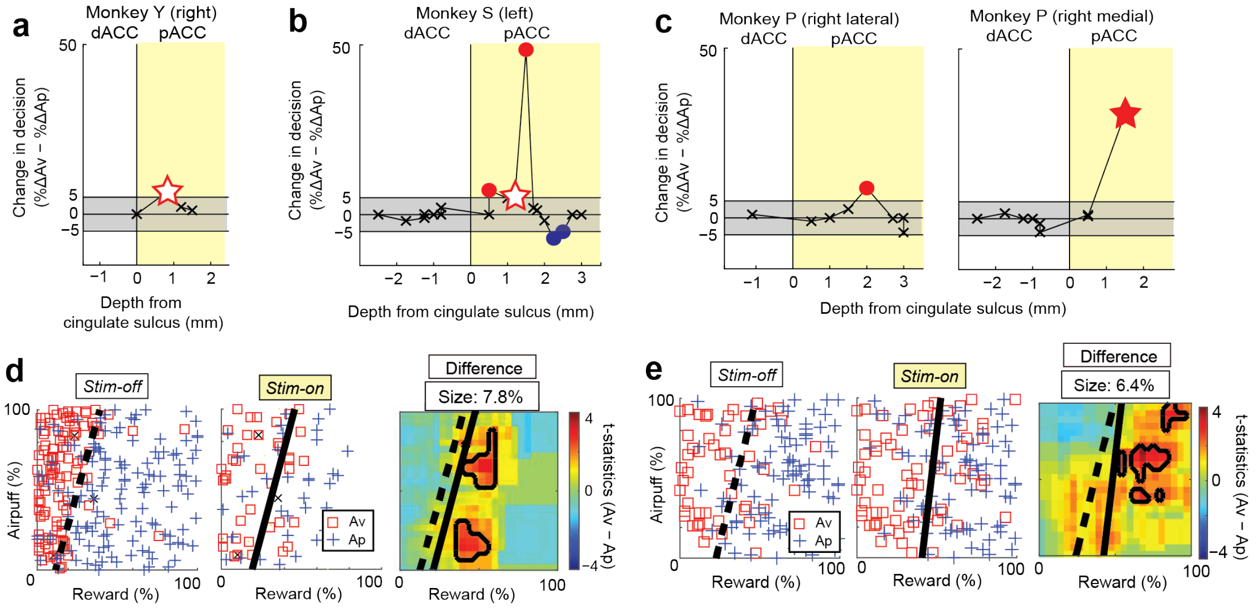


### Supplementary Figure 2

**Microstimulation of the pACC, but not of the dACC, increased avoidance choices in all of three monkeys.**

**a-c)** Changes in decision frequencies (%ΔAv − %ΔAp) by microstimulation in monkeys Y (**a**), S (**b**) and P (**c**), plotted against the depths from the cingulate sulcus. White stars indicate the virus infusion sites where we observed significant increases in Av choices in monkeys Y and S, shown in **d** and **e**, respectively. A red star shows the site where stimulation induced significant changes in decision in monkey P (shown in Fig. 1b) but the tracer virus was not injected. We set a change of 5% as the significance level of the difference in decision matrix between the *Stim-off* and *Stim-on* blocks (see Methods). In each plot, crosses indicate changes below 5%. Red and blue circles indicate the sites where we observed significant increases in the Av and the Ap choices, respectively. **d-e)** Changes in decision frequencies induced by microstimulation at the virus infusion sites in monkeys Y (**d**) and S (**e**), with decision matrices (see Methods) before stimulation (*Stim-off* block, left) and during stimulation (*Stim-on* block, middle), and smoothed differences in decision matrices between *Stim-on* and *Stim-off* blocks (right). Blue crosses indicate Ap choices, and squares indicate Av choices. Dashed and solid lines represent the decision boundary determined by the logistic regression for the *Stim-off* and *Stim-on* blocks, respectively.

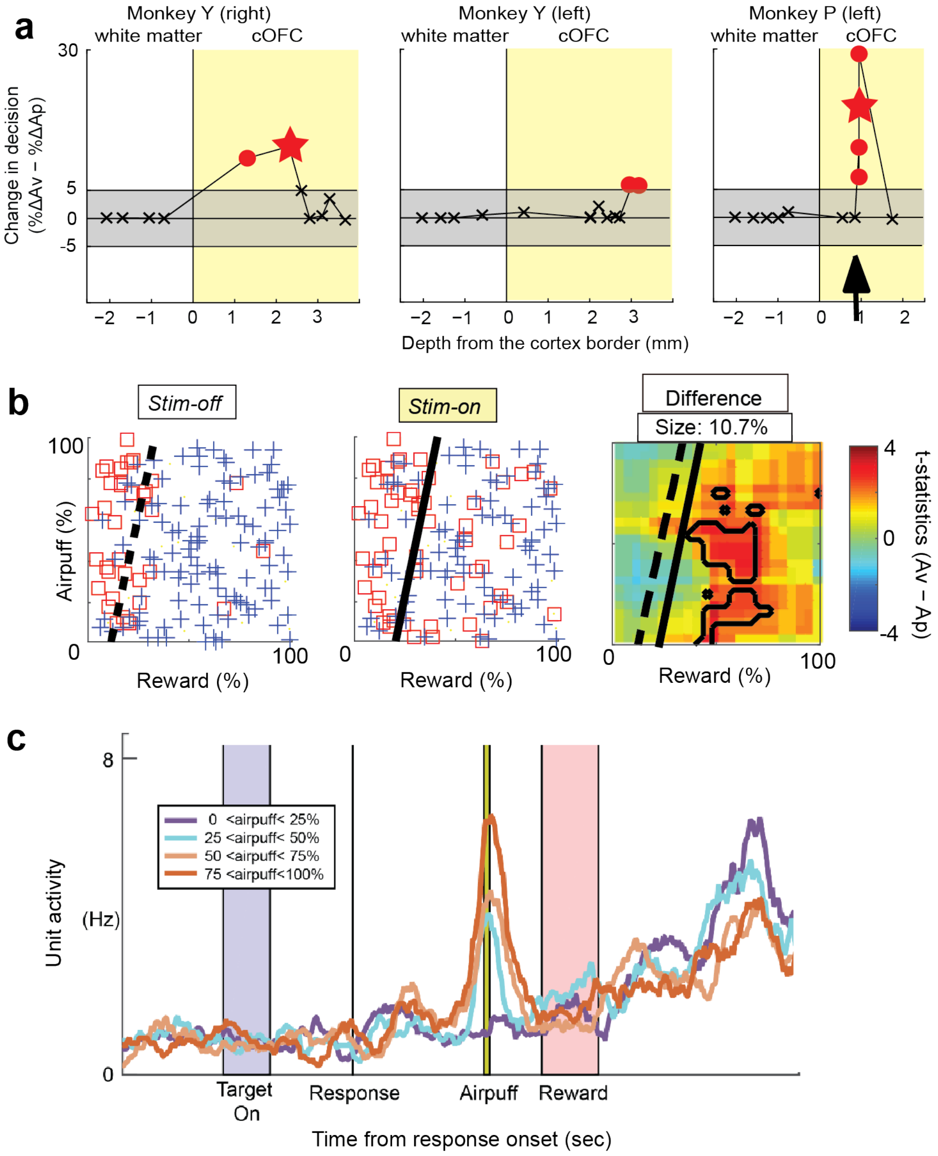


**Supplementary Figure 3**

**Microstimulation of the cOFC, but not surrounding sites, increased Av choices in two monkeys.**

**a)** Changes in decision frequencies (%ΔAv − %ΔAp) by microstimulation in monkeys Y (left and middle panels) and P (right panel), plotted against the depths from the border between white and gray matters. Changes in decision in monkeys Y and P (red stars) are shown in **b** and Fig. 1b, respectively. **b)** Example of the effective stimulation observed in monkey Y (red star in **a**), shown as in Supplementary Fig. 2d. **c)** A cOFC unit that encoded the delivery of an aversive stimulus. This unit was recorded at the site where microstimulation increased Av decisions in monkey P (arrow in **a**). The unit increased its firing rate, expecting delivery of the aversive stimulus (airpuff), and the magnitude of the increase depended on the strength of the airpuff.

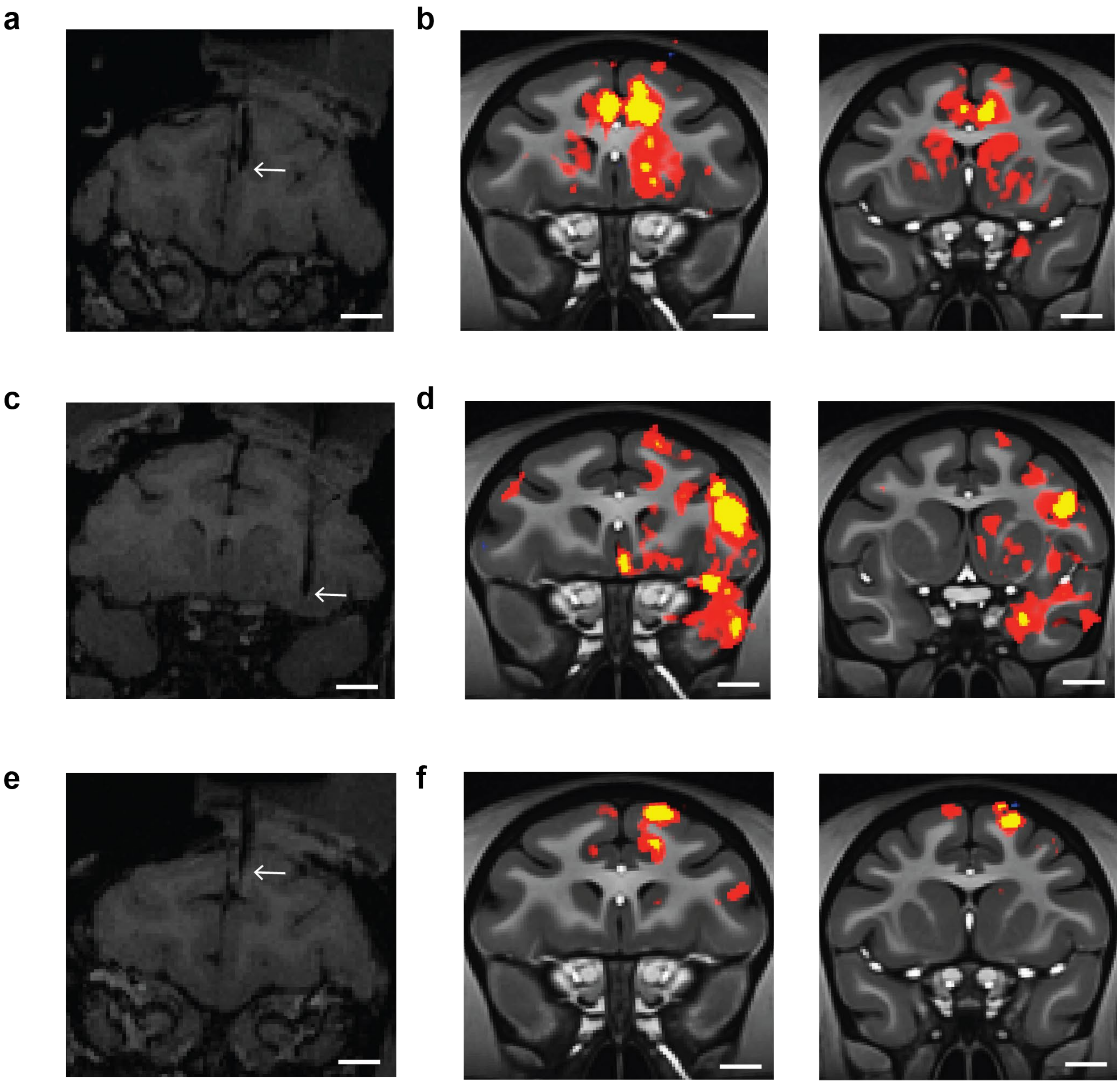


**Supplementary Figure 4**

**Microstimulation of the pACC and cOFC induced striatal activation visualized by fMRI.**

**a)** Location of the stimulation electrode in the pACC (arrow) visualized by MRI in monkey N. **b)** Strong activation induced by pACC microstimulation was detected in the anterior striatum by fMRI. Note that the activation was seen in both hemispheres and was stronger in the anterior to central region, consistent with the histology results. **c)** Location of the stimulation electrode in the cOFC (arrow) visualized by MRI in monkey N. **d)** Strong activation was induced by cOFC microstimulation in the anterior-to-central striatum. Note that the activation was seen only in the right (ipsilateral) hemisphere, in agreement with histology results. **e)** Location of the stimulation electrode in the dorsal ACC (dACC, arrow) visualized by MRI in monkey N. This was a control location for the pACC. The electrode track was the same as the pACC, but the tip was located above the cingulate sulcus. **f)** No activation was found in the striatum following dACC microstimulation. Scale bars represent 5 mm.

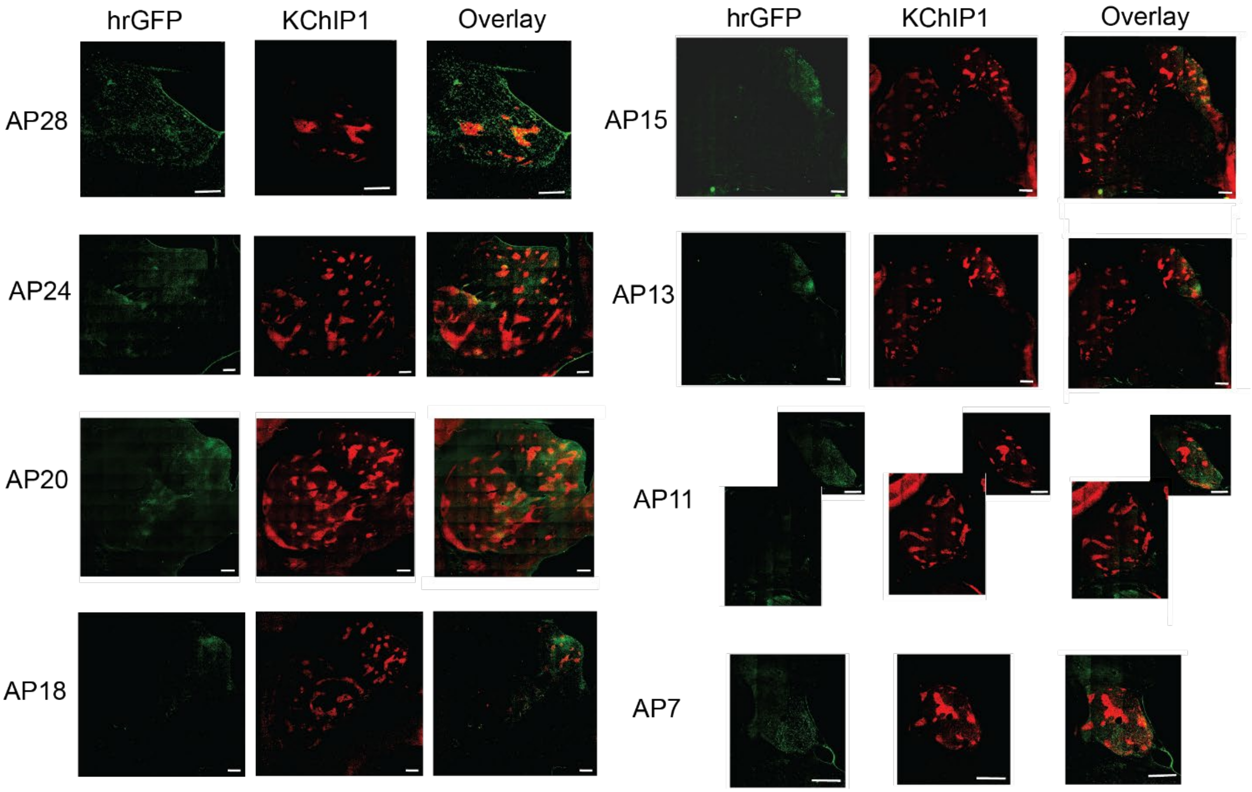


**Supplementary Figure 5**

**The pACC projects preferentially to ipsilateral striosomes.**

Representative images of left striatal sections from monkey S. These sections were double-stained for hrGFP (virus tracer, left) and KChIP1 (striosome marker, middle). Images at right show the overlay of hrGFP and KChIP1 images. Scale bars represent 1 mm.

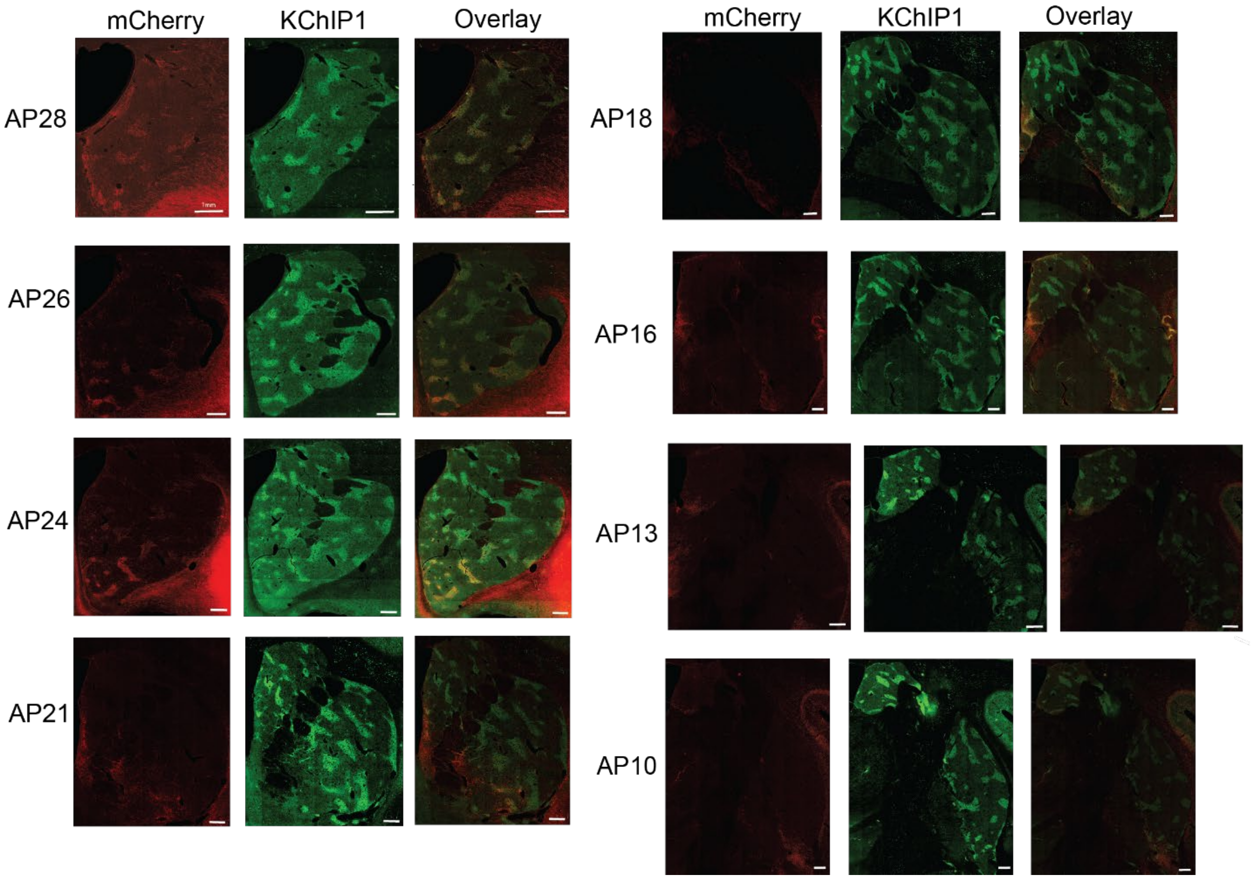


**Supplementary Figure 6**

**The cOFC projects preferentially to ipsilateral striosomes.**

Representative images of right striatal sections from monkey A. These sections were double-stained for mCherry (virus tracer, left) and KChIP1 (striosome marker, middle). Right images show the overlay of mCherry and KChIP1 images. Scale bars show 1 mm.

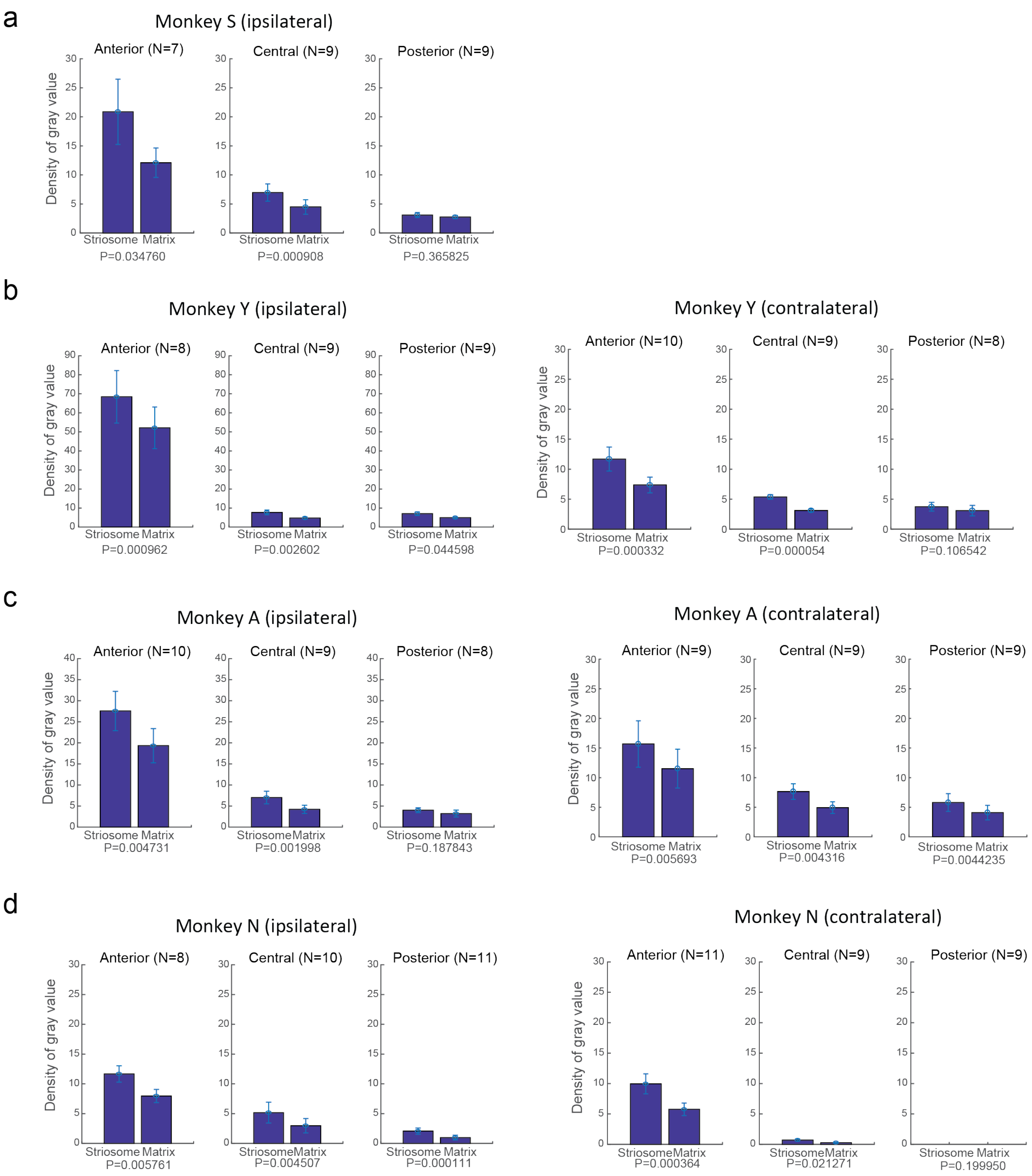


**Supplementary Figure 7**

**Summary of pACC projections to ipsilateral and contralateral striatum.**

Density of projections from the pACC to ipsilateral (left panels) and contralateral (right panels) striatum in monkeys S (**a**), Y (**b**), A (**c**) and N (**d**). Both ipsilateral and contralateral projections were more dense in the anterior part than in the posterior part. However, the overall density was smaller for contralateral projections compared to that for ipsilateral projections. For both ipsilateral and contralateral striatum, pACC projections to striosomes were significantly denser than those to matrix in anterior and middle regions. Gray values (fiber density) were calculated from fluorescent images of histological sections (Methods), and striatal sections were classified into anterior (AP28-22), central (AP21-14) and posterior (AP13-7) regions. Error bars represent the standard error of the mean. The number of sections (N) and the results of paired t-tests (P) are indicated for each plot.

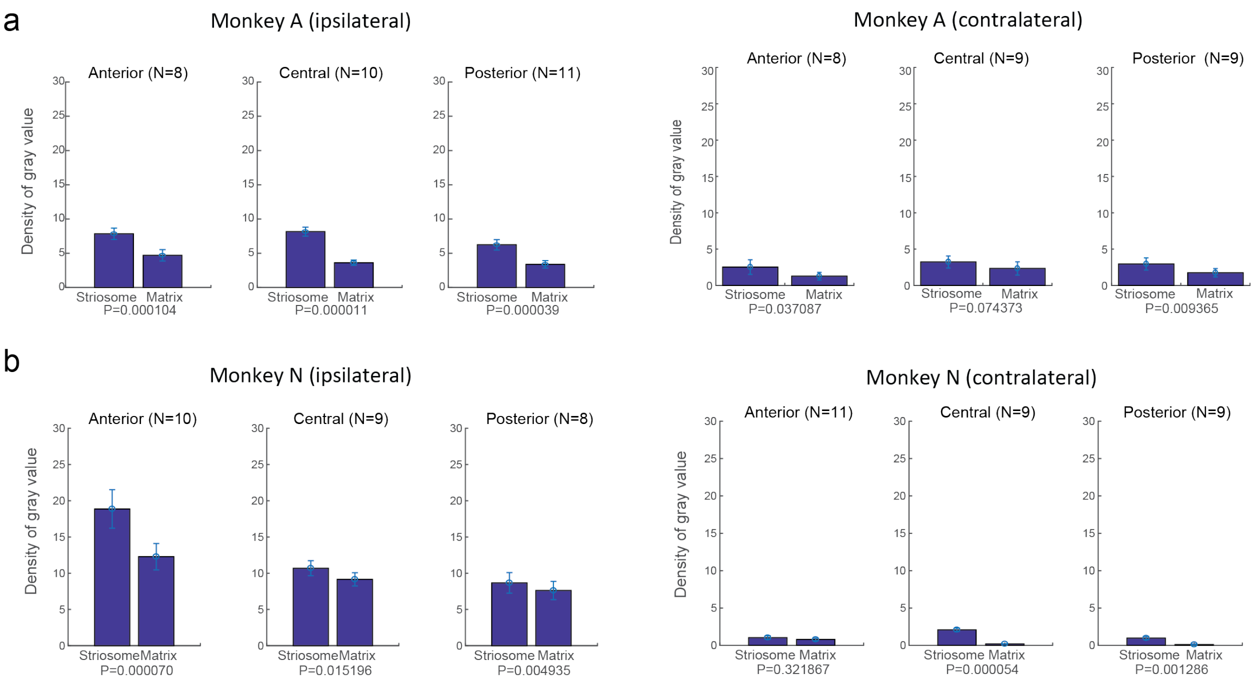


**Supplementary Figure 8**

**Summary of cOFC projects to ipsilateral and contralateral striatum.**

Density of cOFC projections to ipsilateral (right hemisphere) striatum in monkeys A (**a**) and N (**b**), shown as in Supplementary Fig. 7. The cOFC projections to ipsilateral striosomes were significantly greater than those to matrix in all regions in monkey A. Contralateral projections were weak in both monkeys.

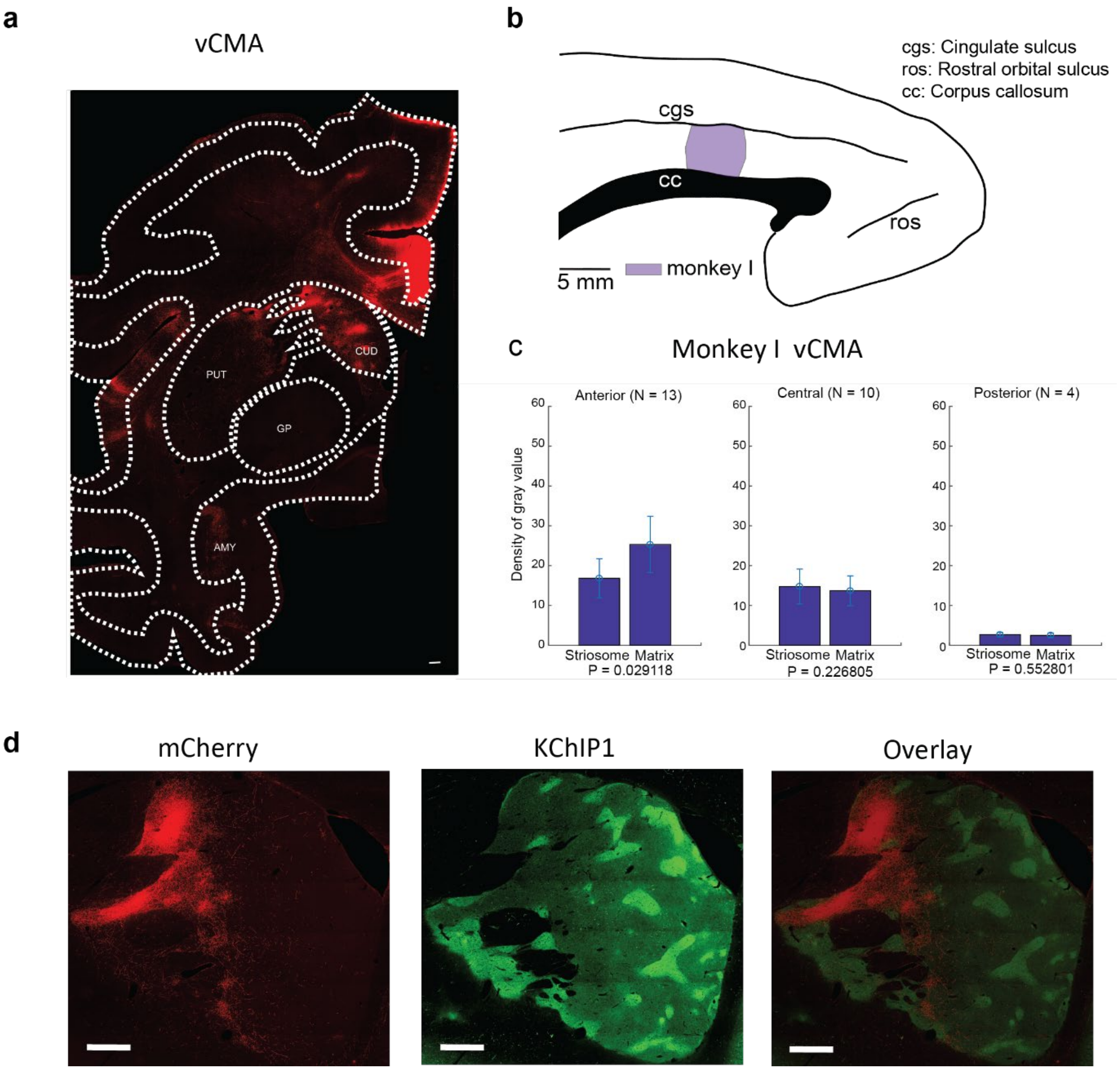


**Supplementary Figure 9**

**Summary of projections from the ventral cingulate motor area (vCMA) to ipsilateral matrix.**

**a)** A sample image of the vCMA injection site in monkey I, stained for mCherry (red). CUD: Caudate, PUT: Putamen, GP: Globus Pallidus, AMY: Amygdala. **b)** Schematic illustration of tracer injection sites (purple), shown in medial views of the left hemisphere. **c)** Summary of vCMA projections to ipsilateral (left hemisphere) striatum in monkey I. Projections to ipsilateral matrix were significantly greater than those to striosomes in the anterior region. **d)** Images of the anterior striatum section of monkey I, shown as in Supplementary Figs. 5 and 7. Scale bars represent 1 mm.
