## Supplementary Table 1 for "Microstimulation of primate neocortex targeting striosomes induces negative decision-making"

**Supplementary Table 1.** Summary of monkeys used in this study and the experimental procedures applied to each monkey. L: left hemisphere, R: right hemisphere. F: female, M: male. Data from the left hemisphere of monkey Y with cOFC injection were not included due to the leakage of virus tracer to other cortical regions.

| Monkey name (sex) | Stimulation during ApAv | Recording during ApAv | fMRI with stimulation | Virus injection (total amount) |
|---|---|---|---|---|
| Monkey S (F) | L & R pACC | R pACC | No | AAVDJ-CMV-hrGFP in LpACC (0.9 μl) |
| Monkey P (F) | LpACC, and LcOFC | LpACC, and LcOFC | No | No |
| Monkey A (F) | No | No | No | AAVDJ-CMV-hrGFP in RpACC (1.5 μl)<br>AAVDJ-CMV-mCherry in RcOFC (1 μl) |
| Monkey Y (F) | RpACC, and L & R cOFC | No | No | AAVDJ-CMV-mCherry in RpACC (2 μl)<br>*AAVDJ-CMV-hrGFP in LcOFC (1.4 μl) *Data not included |
| Monkey N (M) | No | No | RpACC, and RcOFC | AAVDJ-CMV-mCherry in RpACC (1 μl)<br>AAVDJ-CMV-hrGFP in RcOFC (0.9 μl) |
| Monkey I (M) | No | No | No | AAVDJ-CMV-mCherry in LvCMA (1.4 μl) |
